# NMDA Receptor Subunit NMR-2 Regulates Pathogen-Induced Immune Responses via the Nervous System in *C. elegans*

**DOI:** 10.64898/2026.05.29.728750

**Authors:** Benson Otarigho, Jonathan Lalsiamthara, Alejandro Aballay

**Affiliations:** Department of Genetics, The University of Texas MD Anderson Cancer Center, Houston, TX; Department of Microbiology and Molecular Genetics, McGovern Medical School at UTHealth, Houston, TX

**Keywords:** Glutamate, Innate Immunity, NMR-2, Pathogen resistance, Sensory neurons

## Abstract

Neural control of innate immunity must balance restraint of basal immune activity with rapid activation upon pathogen encounter. Glutamate, the primary excitatory neurotransmitter in the nervous system, has been implicated in several neurological disorders associated with inflammation, suggesting a potential link to immune regulation. However, how glutamatergic signaling contributes to immune balance remains unknown. Here, we demonstrated that the NMDA-type ionotropic glutamate receptor subunit NMR-2, a component of the NMDA receptor complex, acts in the *C. elegans* nervous system as a key regulator of pathogen-induced immune responses. Loss of *nmr-2* enhanced resistance to *Staphylococcus aureus* and *Pseudomonas aeruginosa*, without elevating basal immune gene expression. The pathogen-induced response controlled by NMR-2 required the conserved PMK-1/p38 MAPK, DAF-16/FOXO, and HLH-30/TFEB pathways. We identified the AVD interneuron as the site of action of NMR-2, where it integrates inputs from upstream sensory neurons ASE, ASK, AQR, and PQR. These findings uncover a neural circuit in which glutamatergic signaling distinguishes basal immune gene expression from pathogen-induced immune responses, revealing a mechanism that promotes effective defense without altering baseline immunity.

## Introduction

Activation of immune responses must be tightly controlled to enable rapid, robust responses upon pathogen encounter while preventing the deleterious effects of unrestrained activity ^1, 2^. Excessive basal immune activity imposes metabolic and reproductive costs, interferes with growth, and may desensitize inducible defense programs. Conversely, insufficient immune response compromises survival in the presence of pathogens ^3, 4, 5^. Neural circuits are well-positioned to balance these opposing requirements, as they integrate sensory inputs with internal state to modulate systemic physiology ^6, 7, 8^. Work in *Caenorhabditis elegans* has established that the nervous system exerts extensive control over immunity by releasing neurotransmitters and neuromodulators that act on intestinal and systemic defense pathways ^8, 9, 10, 11, 12^. Identified pathways include serotonergic, dopaminergic, octopaminergic, and GABAergic signals that primarily tune basal immune tone, either suppressing or elevating effector gene expression even in the absence of pathogens ^7, 8, 9, 10, 11, 12, 13, 14, 15, 16^. What has remained unclear is whether the nervous system also encodes a dedicated circuit that specifically controls pathogen-induced activation of immunity while retaining basal activity.

Glutamatergic signaling is a major excitatory modality in vertebrate and invertebrate nervous systems. Ionotropic NMDA, AMPA, and kainate receptors mediate fast synaptic transmission, whereas metabotropic receptors support slower modulation ^17, 18, 19, 20^. In *C. elegans*, NMDA receptors are heteromeric complexes containing the essential subunit NMR-1 together with the modulatory subunit NMR-2. These receptors are prominent in interneurons involved in sensorimotor integration and the state-dependent control of behavior ^21, 22, 23, 24^. Despite this central function, the contribution of glutamate receptors to host defense in *C. elegans* has not been defined.

In *C. elegans*, neural control of intestinal defense converges on conserved transcriptional regulators, including PMK-1 p38 MAPK, DAF-16 FOXO, and HLH-30 TFEB, which coordinate stress and immune effectors ^25, 26, 27^. Prior studies have mapped several neurotransmitter pathways that significantly set basal immune tone. Octopaminergic inputs regulate intestinal protein synthesis and suppress defense gene expression ^8, 14, 15, 16^, GABAergic programs coordinate immunity with longevity outputs ^10, 12^, and serotonergic and dopaminergic cues adapt defense to environmental context ^13^. Interneurons that receive convergent sensory input and broadcast command signals provide an anatomical substrate for state-dependent control of systemic physiology ^28, 29^. Among receptors expressed in these cells, NMDA-type glutamate receptors are well suited to gate inducible responses because their activation requires coincident presynaptic glutamate release and postsynaptic depolarization, a property that restrains basal activity yet allows strong activation when inputs align ^17, 21, 22, 30^. This synaptic integration mode contrasts with diffuse neuromodulatory signaling, providing a mechanistic basis to test whether glutamatergic transmission gates pathogen-elicited immunity while limiting unnecessary basal activation.

We therefore asked whether glutamate receptor genes in *C. elegans* control the immune state in a way that separates basal repression from pathogen-elicited activation. We focused on glutamate receptors because their subunit composition provides a mechanism for modulating synaptic gain and shaping temporal integration in interneurons that link sensory inputs to downstream command layers ^30, 31, 32^. Using genetic analysis of glutamate receptor mutants, transcriptional profiling under basal and infected conditions, and placement into defined circuits by neuronal rescue and ablation, we tested whether glutamatergic signaling links sensory context to intestinal defense through canonical pathways.

We found that loss of the NMDA receptor subunit NMR-2 confers significant pathogen resistance to multiple bacterial pathogens, including *Staphylococcus aureus* and *Pseudomonas aeruginosa*. We further demonstrated that pathogen resistance is associated with a pathogen-induced immune response, characterized by the upregulation of immune effector genes that are known to be under the regulation of the PMK-1, DAF-16, and HLH-30 pathways. This suggests that NMR-2 differs from previously characterized neurotransmitter receptors in selectively influencing robust pathogen-induced immune responses. Further experiments indicate that NMR-2 mediates this pathogen-induced immune regulation through the AVD interneuron, which functions within a defined glutamatergic neural circuit. The immune role of NMR-2 through AVD neurons may be mediated by the individual or combined contributions of upstream sensory neurons such as ASE, ASK, and AQR/PQR. Our findings reveal a previously unrecognized role for glutamatergic signaling in enabling robust pathogen-induced responses without altering basal immune gene expression, with NMR-2 acting as a key neural modulator that limits harmful immune overactivation and highlights conserved immune mechanisms.

## Results

### Loss of function of the NMDA-type ionotropic glutamate receptor subunit NMR-2 enhances pathogen resistance

To explore the mechanisms underlying pathogen-induced immune responses, we reviewed prior studies on neurotransmitter signaling, particularly receptors, previously implicated in the control of immunity in *C. elegans* (Supplementary Table 1). These studies, together with our analysis, indicate that several neurotransmitter receptors modulate basal immune activity, either by enhancing or suppressing it. Notably, glutamatergic signaling has remained largely unexplored in this context. Therefore, we conducted a targeted genetic screen of mutants in genes encoding glutamate receptors in *C. elegans,* which includes ionotropic NMDA, AMPA, and kainate as well as metabotropic receptors (Figure 1A). We found that animals lacking *nmr-2* [*nmr-2(ok3324)*], which encodes a regulatory subunit of the NMDA-type ionotropic glutamate receptor, survived significantly longer than wild type on a full lawn of *Staphylococcus aureus* NCTCB325 across independent experiments and plates (Figure 1B–D). In contrast, mutants in other glutamate receptor genes (iGluR and mGluR) showed survival rates comparable to wild-type animals (Figure 1B–D). To test whether *nmr-2* itself mediates pathogen resistance, we expressed *nmr-2* under its endogenous promoter, which rescued the pathogen-resistant phenotype of *nmr-2(ok3324)* animals in survival assays (Figure S1A). We further examined whether pharmacological inhibition of NMDA receptor signaling affects pathogen resistance using dextromethorphan (DXM). WT and *nmr-2(ok3324)* animals treated with DXM were exposed to a full lawn of *S. aureus* (Figure S1B). DXM treatment significantly enhanced survival in WT animals, whereas it had no additional effect on *nmr-2(ok3324)* mutants, consistent with both perturbations acting through the same NMDA receptor-dependent pathway ^33, 34^. These findings indicate that the regulatory and signaling roles of NMR-2 subunits are critical in modulating host-pathogen interactions in *C. elegans*. Although DXM is widely used as an NMDA receptor antagonist, it is not fully selective and may exhibit off-target effects ^33, 34^. Therefore, the pharmacological results should be interpreted cautiously and are presented as supportive, rather than definitive, evidence for NMR-2-dependent glutamatergic signaling in immune regulation.

**Figure 1.**
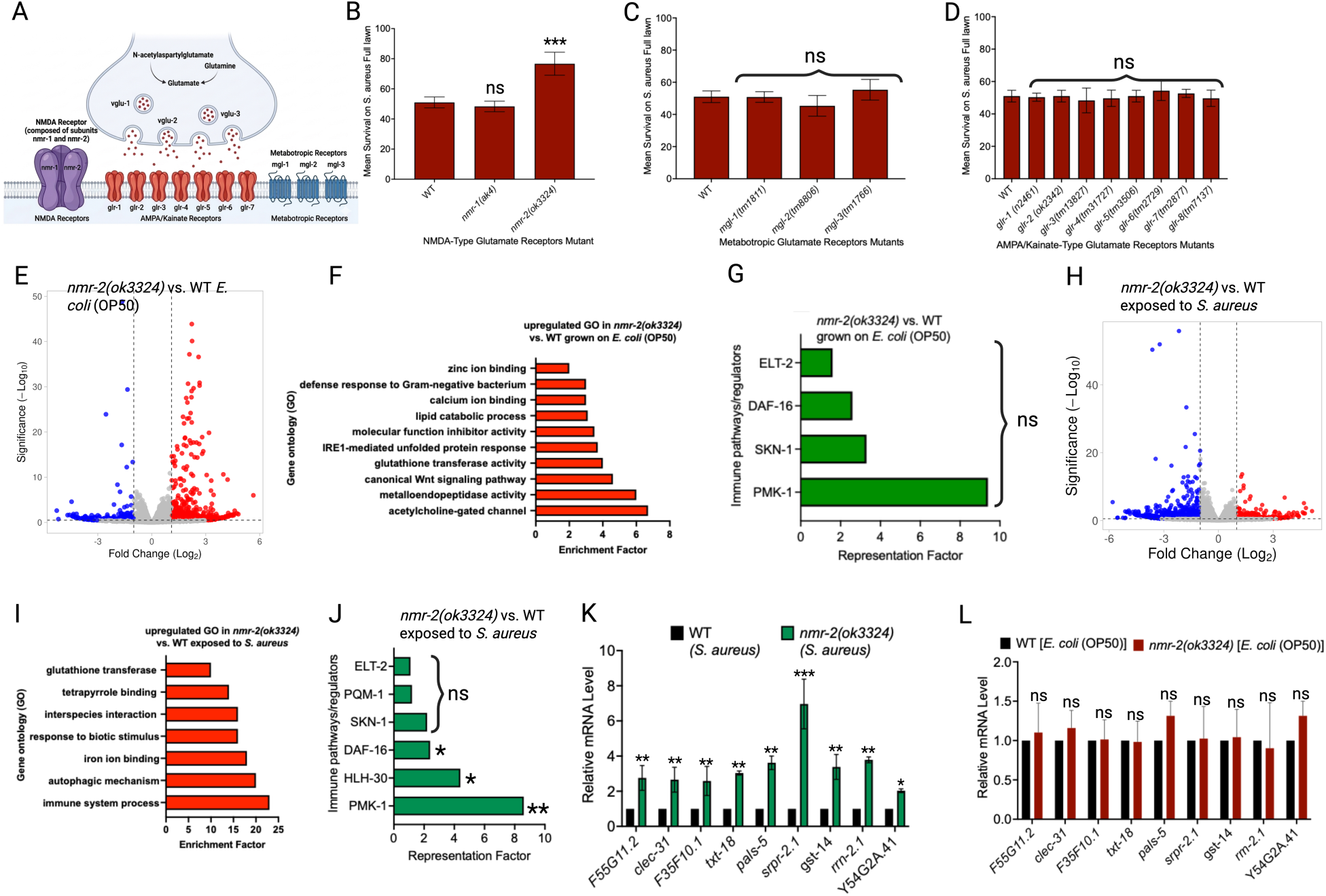
A mutation in Glutamate Receptor, NMR-2, exhibits pathogen-induced upregulation of immune genes. A. *C. elegans* putative glutamatergic receptors showing Ionotropic NMDA, AMPA, and kainate as well as metabotropic receptors. B. WT and mutants of the NMDA-type glutamate receptor were exposed to a full lawn of *S. aureus*, and their mean survival was measured. Bars represent mean values, while error bars indicate SD; NS, Not Significant; *** P < 0.0001. C. WT and mutants of the metabotropic glutamate receptor were exposed to a full lawn of *S. aureus*, and their mean survival was measured. D. WT and mutants of the AMPA/Kainate-type glutamate receptor were exposed to a full lawn of *S. aureus*, and their mean survival was measured. E. Volcano plot of upregulated and downregulated genes in *nmr-2(ok3324)* versus WT animals grown on *E. coli* (OP50). Red and blue dots represent significantly upregulated and downregulated genes, respectively, while the gray dots represent non-significant genes. F. Gene ontology analysis of upregulated genes in *nmr-2(ok3324)* versus WT animals grown on *E. coli* (OP50). The results were filtered to include only significantly enriched terms with a q-value < 0.1, with minor modifications made by merging similar Gene Ontology terms where appropriate. G. Representation factors of immune pathways for the upregulated immune genes in *nmr-2(ok3324)* versus WT animals grown on *E. coli* (OP50). Each bar denotes a representation factor. Statistical significance is denoted as NS, not significant, *p < 0.05, and **p < 0.001. H. Volcano plot of upregulated and downregulated genes in *nmr-2(ok3324)* versus WT animals exposed to *S. aureus* for four hours. Red and blue dots represent significantly upregulated and downregulated genes, respectively, while the gray dots represent non-significant genes. I. Gene ontology analysis of upregulated genes in *nmr-2(ok3324)* versus WT animals exposed to *S. aureus* for four hours. The results were filtered to include only significantly enriched terms with a q-value < 0.1, with minor modifications made by merging similar Gene Ontology terms where appropriate. J. Representation factors of immune pathways for the upregulated immune genes in *nmr-2(ok3324)* versus WT animals exposed to *S. aureus* for four hours. Each bar denotes a representation factor. Statistical significance is denoted as NS, not significant, *p < 0.05, and **p < 0.001. K. Quantitative reverse transcription-PCR (qRT-PCR) analysis was performed to examine DAF-16- and PMK-1-dependent immune gene expression in WT and *nmr-2(ok3324)* animals following infection with *S. aureus*. Bars represent mean values, and error bars indicate the standard deviation (SD) from three independent experiments. Statistical significance is denoted as *p < 0.05, **p < 0.001, and ***p < 0.0001. L. Quantitative reverse transcription-PCR (qRT-PCR) analysis was performed to examine DAF-16- and PMK-1-dependent immune gene expression in WT and nmr-2(ok3324) animals fed with *E. coli* OP50. Bars represent mean values, and error bars indicate the standard deviation (SD) from three independent experiments. Statistical significance is denoted as *p < 0.05, **p < 0.001, and ***p < 0.0001.

### NMR-2 mutant displays no significant basal immunity but enhanced pathogen-induced immunity

Since several neurotransmitter receptors linked to immunity in *C. elegans* strongly modulate basal immune tone (Supplementary Table 1), we next asked whether the glutamate receptor *nmr-2*, whose role in immunity is unknown, contributes specifically to basal or pathogen-induced immune responses. We profiled transcriptional outputs under nonpathogenic conditions (live *Escherichia coli* OP50 food) and following *S. aureus* exposure (Figure 1E-J; S2; Supplementary Table 2-3). RNA-seq analysis revealed no significant enrichment of immune-related GO categories at baseline in *nmr-2(ok3324)* relative to wild type, despite modest changes in the expression of some immune genes (Figure 1F-G; Supplementary Table 2). In contrast, infection elicited significant upregulation of defense genes in *nmr-2(ok3324)* compared with wild type (Figure 1I-J; Supplementary Table 3), with PMK-1, DAF-16, and HLH-30-dependent immune genes significantly enriched among the upregulated immune genes (Figure 1J; Supplementary Table 3). The absence of significant changes in basal immune genes, as well as the pathogen-induced upregulation of immune genes, was further confirmed in *nmr-2(ok3324)* animals by quantitative reverse transcription PCR (qRT-PCR) (Figure 1K-L). Together, these analyses indicate that *nmr-2(ok3324)* mutants do not exhibit coordinated immune activation under basal conditions but display selective induction of immune effectors upon infection.

To further assess whether NMR-2 modulates basal immune responses among neurotransmitter receptors, we analyzed published data on octopamine, GABA, dopamine, acetylcholine, and serotonin receptors, which have been reported to alter basal immune gene expression and exhibit additional induction upon infection ^8, 12, 14, 35^. Given the availability of transcriptomic data, we focused on the octopamine receptor OCTR-1, which is well established as a regulator of immune defense against diverse pathogens ^8, 14^. Reanalysis of transcriptomics data from previous studies ^8, 16^ showed that *octr-1(ok371)* animals exhibited upregulation of immune genes under both basal and pathogen-exposed conditions (Figure 2A–D, Supplementary Table 1 & 4), as also demonstrated by Styer, *et al.* (2008). In line with the role of OCTR-1, analysis of previous gene expression data suggests that other neurotransmitter receptors, including GAR-2/3 (acetylcholine), DOP-4 (dopamine), UNC-49 (GABA), and SER-7 (serotonin), also play significant roles in regulating immune responses under both basal and pathogen-exposed conditions (Figure 2E) ^8, 12, 14, 35^. These findings indicate that these receptors broadly regulate immune gene expression under basal and pathogen-exposed conditions, with some receptors suppressing basal immune tone and others promoting immune activation, rather than specifically controlling pathogen-induced responses. Notably, many of the immune genes upregulated under basal conditions overlap with those induced upon infection, suggesting that these receptors primarily influence basal immune tone rather than selectively controlling inducible immune responses. In contrast, our findings indicate that the NMDA receptor subunit NMR-2 selectively modulates pathogen-induced immune responses without broadly affecting basal immune gene expression, allowing a robust response during infection. Thus, uncovering a neuronal mechanism that accounts for the excessive basal activation of innate immunity suggests that such hyperactivity may compromise defense by limiting inducibility or imposing physiological costs, ultimately heightening susceptibility to infection ^3, 4, 5, 36^. To confirm these patterns at the single-animal level, we used strain AY101 [*pirg-5p::GFP* + *rol-6(su1006)*], which reports intestinal *irg-5* expression with a large dynamic range. Under *E. coli* conditions, AY101 animals with upregulated basal immune genes displayed bright GFP, whereas those with low or undetectable basal expression showed little GFP. Consistently, animals with strong GFP signals were more susceptible to *S. aureus* infection than those lacking detectable GFP (Figure S3A). We further performed qRT-PCR analysis to quantify immune gene expression in animals with high and low GFP levels, confirming that bright GFP animals exhibit higher expression of immune effector genes compared to those with low or undetectable GFP (Figure S3B). Together, these findings indicate that, unlike other neurotransmitter receptors that broadly modulate basal immune gene expression, NMR-2 selectively influences pathogen-induced immune responses without significantly affecting basal immune activity.

**Figure 2.**
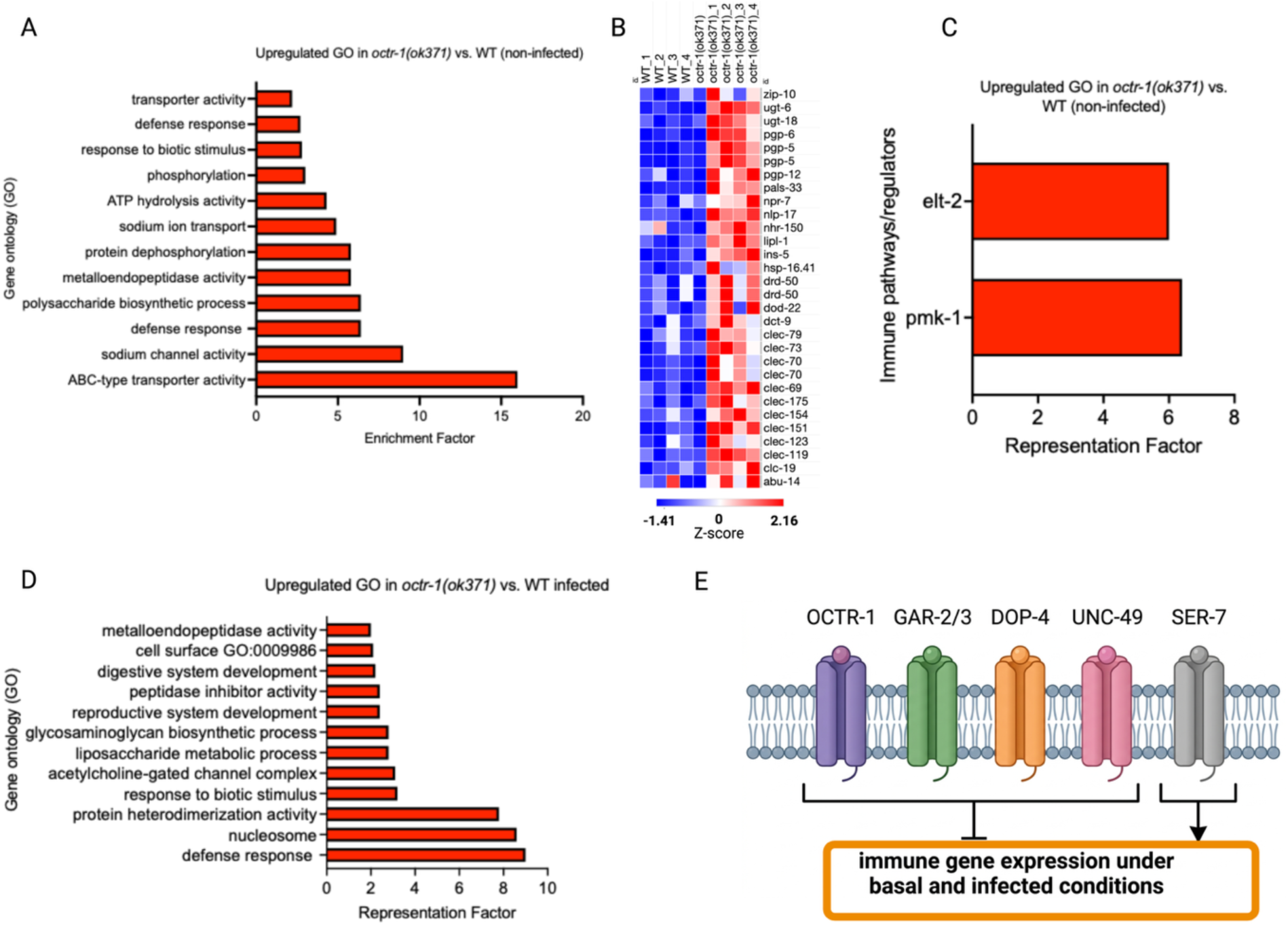
Neurotransmitter receptors previously implicated in immunity broadly regulate immune gene expression under basal and pathogen-exposed conditions rather than selectively controlling pathogen-induced responses. A. Gene ontology analysis of upregulated genes in *octr-1(ok371)* versus WT animals that were not exposed to a pathogen. The results were filtered to include only significantly enriched terms with a q-value < 0.1, with minor modifications made by merging similar Gene Ontology terms where appropriate. B. Heatmap of the most upregulated immune genes in *octr-1(ok371)* versus WT animals that were not exposed to a pathogen. C. Representation factors of immune pathways for the upregulated immune genes in *octr-1(ok371)* versus WT animals. D. Gene ontology analysis of upregulated genes in *octr-1(ok371)* versus WT animals that were exposed to a pathogen. The results were filtered to include only significantly enriched terms with a q-value < 0.1, with minor modifications made by merging similar Gene Ontology terms where appropriate. E. A schematic representation of different neurotransmitter receptors known to either suppress or activate immune gene expression under both basal and pathogen-infected conditions.

### The enhanced survival of *nmr-2* mutants driven by pathogen-induced immunity is independent of bacterial colonization or physiological changes

We tested whether the survival phenotype generalizes beyond a single pathogen. *nmr-2(ok3324)* animals survived longer than wild type on *Pseudomonas aeruginosa* PA14 (Figure 3A–B) and showed no survival advantage on nonpathogenic live or UV-killed *E. coli* OP50 (Figure 3C–D), indicating that the phenotype reflects infection outcomes rather than pathogen-independent longevity effects. Since it is known that altered pathogen load can confound survival differences ^10, 37, 38^, we quantified intestinal accumulation of GFP-labeled *P. aeruginosa* and performed colony-forming unit counts 24 hours after exposure. Bacterial burden did not differ significantly between *nmr-2(ok3324)* and wild type (Figure 3E–F). However, CFU quantification 24 hours after *S. aureus* exposure revealed that *nmr-2(ok3324)* animals harbor a higher bacterial load compared to wild-type animals (Figure 3G). Thus, improved survival in *nmr-2(ok3324)* animals is not explained by reduced colonization in our assays, indicating that enhanced survival reflects increased host tolerance rather than improved pathogen clearance. We also evaluated behaviors and physiological parameters known to affect infection outcomes. Pathogen lawn occupancy (avoidance), pharyngeal pumping following pathogen exposure, defecation cycle timing, and brood size were comparable between *nmr-2(ok3324)* and wild type (Figure S4A–D). Given the influence of avoidance, feeding, defecation, and reproduction on fitness and survival, the absence of differences across these measures supports an immune-centric explanation for the *nmr- 2* survival phenotypes under the tested conditions.

**Figure 3.**
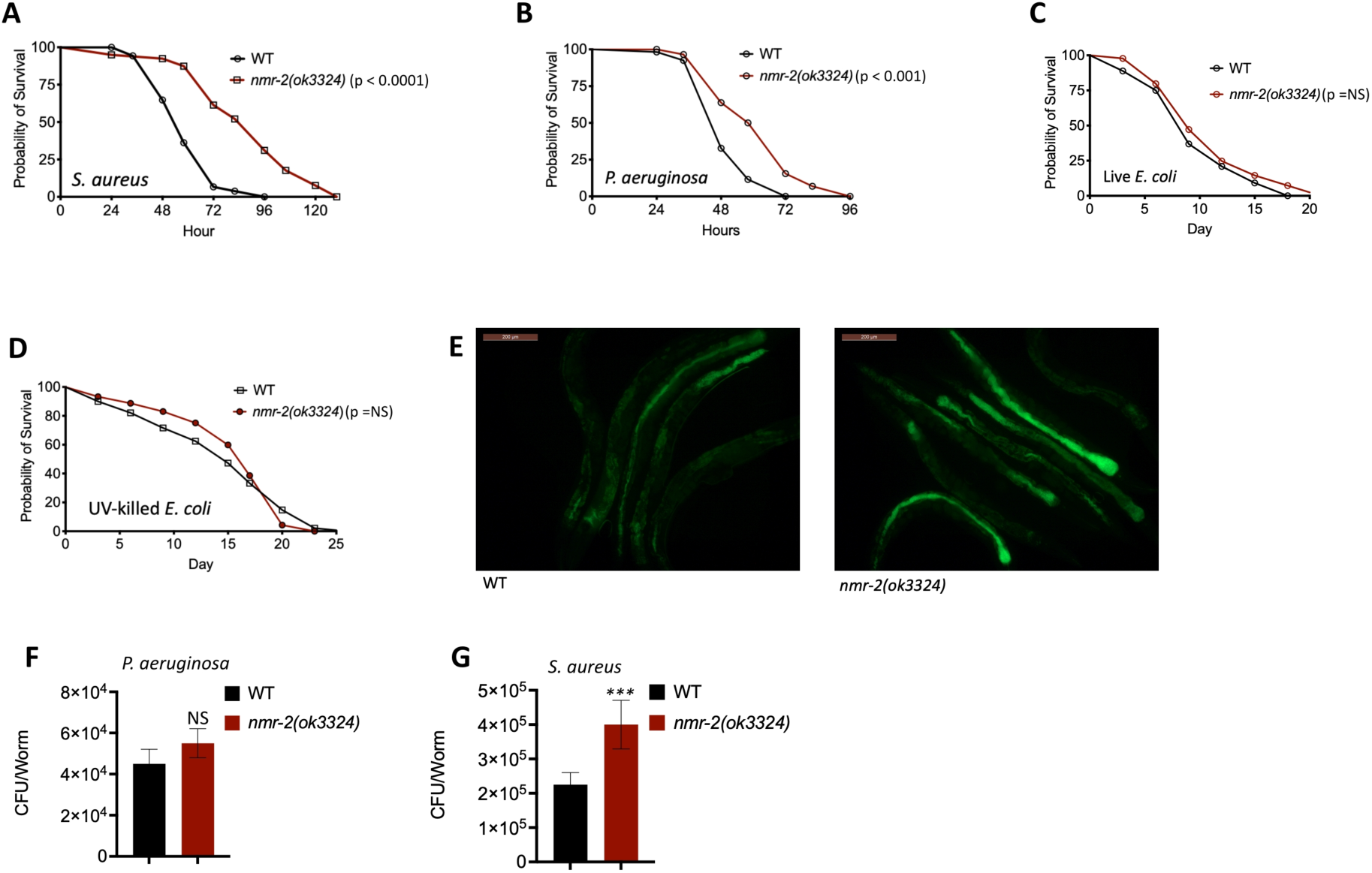
A mutation in the glutamate receptor NMR-2 enhances pathogen resistance through pathogen-induced immune responses when animals are exposed to different pathogens. A. WT, *nmr-2(ok3324)* were exposed to S. aureus full lawn and scored for survival. B. WT, *nmr-2(ok3324)* were exposed to *P. aeruginosa* full lawn and scored for survival. C. WT, *nmr-2(ok3324)* on live *E. coli* (OP50) scored for survival. D. WT, *nmr-2(ok3324)* on UV-killed *E. coli* (OP50) scored for survival. E. Colonization of WT, and *nmr-2(ok3324)* animals by *P. aeruginosa*-GFP after 24 h at 25°C (n = 10). Scale bar, 200 μm. F. Colony-forming units per WT and *nmr-2(ok3324)*, grown on *P. aeruginosa* for 24 h at 25°C (n = 10). Bars represent mean values, and error bars indicate the standard deviation (SD) from three independent experiments. Statistical significance is denoted as NS, not significant. Ten animals were used for each condition. G. Colony-forming units per WT and *nmr-2(ok3324)*, grown on *S. aureus* for 24 h at 25°C (n = 10). Bars represent mean values, and error bars indicate the standard deviation (SD) from three independent experiments. Statistical significance is denoted as ***, significant (p > 0.0001). Ten animals were used for each condition.

### NMR-2-dependent immune response requires PMK-1, DAF-16, and HLH-30 signaling pathways

Following the upregulation of immune genes and regulators mentioned earlier from the transcriptional changes (Figure 1I), we further aimed to uncover the molecular pathways through which NMR-2 modulates immunity. We conducted genetic epistasis analyses using RNAi to knock down key immune regulators in the *nmr-2(ok3324)* animals. The pathogen resistance phenotype exhibited by *nmr-2(ok3324)* animals was partially suppressed by RNAi-mediated knockdown of *daf-16, pmk-1*, or *hlh-30* individually (Figure 4A–C). Simultaneous knockdown of all three immune regulators fully suppressed the survival phenotype (Figure 4D). However, knockdown of other immune-associated transcription factors, such as *skn-1*, did not suppress the survival phenotype exhibited by *nmr-2(ok3324)* animals (Figure 4E). The RNAi that suppressed the *nmr-2(ok3324)* phenotype (Figure 4A-C) was further validated using mutants for the corresponding immune regulators (Figure 4F-H). These findings suggest that NMR-2-dependent pathogen-induced immune responses require immune gene expression associated with the PMK-1, DAF-16, and HLH-30 pathways, providing insight into how a neuronal glutamate receptor influences intestinal transcriptional responses to pathogens.

**Figure 4.**
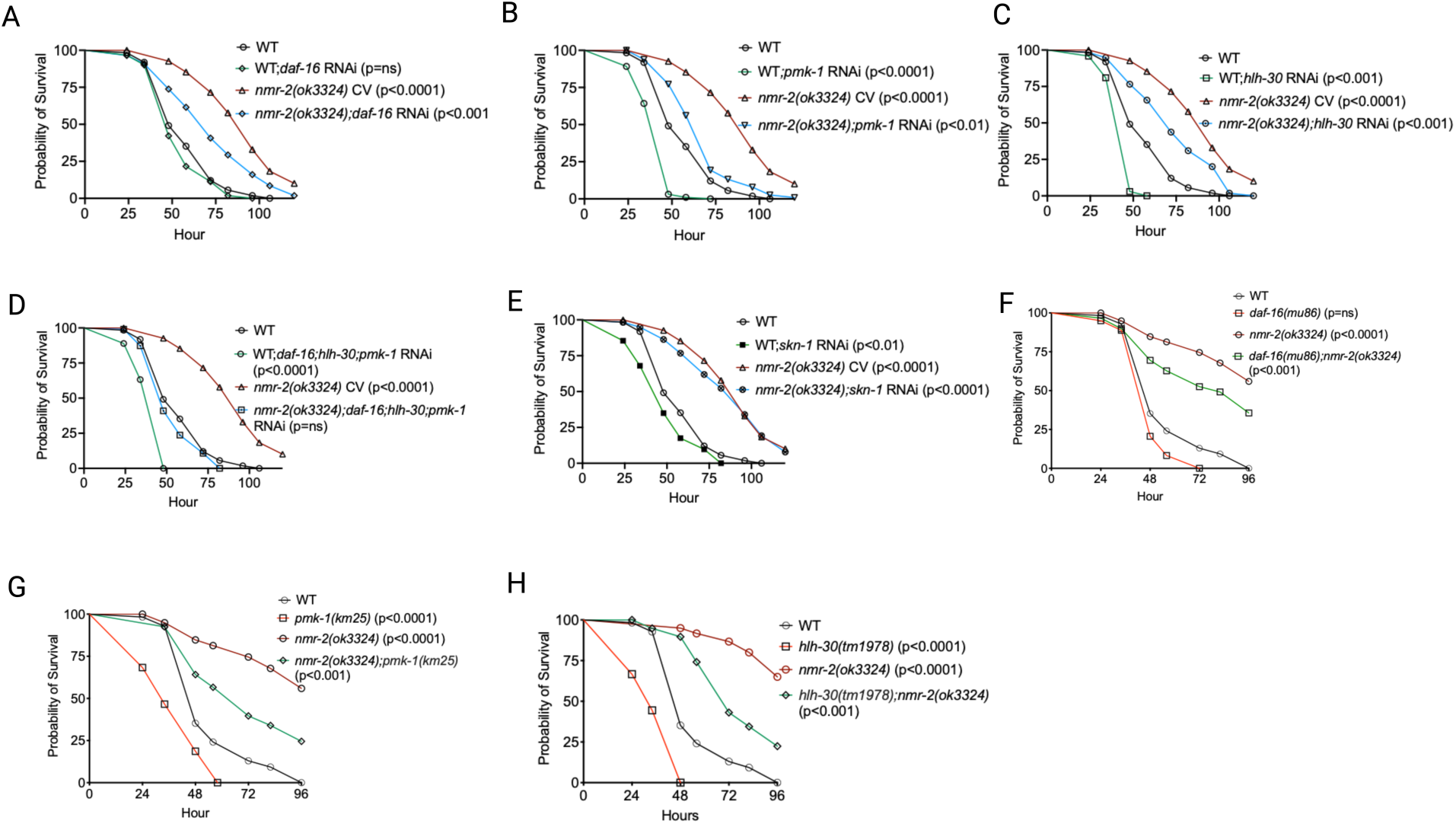
Functional loss of the glutamate receptor NMR-2 promotes immune responses via the DAF-16, PMK-1, and HLH-30 signaling pathways. A. WT, *daf-16* RNAi, *nmr-2(ok3324)*, and *nmr-2(ok3324)*;*daf-16* RNAi animals were exposed to *S. aureus* and scored for survival. B. WT, *pmk-1* RNAi, *nmr-2(ok3324)*, and *nmr-2(ok3324)*;*pmk-1* RNAi animals were exposed to *S. aureus* and scored for survival. C. WT, *hlh-30* RNAi, *nmr-2(ok3324)*, and *nmr-2(ok3324)*;*hlh-30* RNAi animals were exposed to *S. aureus* and scored for survival. EV, empty vector RNAi control. D. WT and *nmr-2(ok3324)* were grown on *daf-16, hlh-30*, and *pmk-1*; RNAi and exposed to *S. aureus* and scored for survival. EV, empty vector RNAi control. E. WT, *skn-1* RNAi, *nmr-2(ok3324)*, and *nmr-2(ok3324)*;*skn-1* RNAi animals were exposed to *S. aureus* and scored for survival. EV, empty vector RNAi control. F. WT, *daf-16(mu86)*, *nmr-2(ok3324)*, and *daf-16(mu86);nmr-2(ok3324)* animals were exposed to *S. aureus* and scored for survival. G. WT, *pmk-1(km25)*, *nmr-2(ok3324)*, and *nmr-2(ok3324);pmk-1(km25)* animals were exposed to *S. aureus* and scored for survival. H. WT, *hlh-30(tm1978)*, *nmr-2(ok3324)*, and *hlh-30(tm1978);nmr-2(ok3324)* animals were exposed to *S. aureus* and scored for survival.

### NMR-2 Regulates Pathogen-Induced Immune Response via the AVD Interneuron

Given that NMR-2 is primarily expressed in the nervous system ^22^, we first expressed *nmr-2* under the pan-neuronal promoter *Prab-3.* This rescued the enhanced survival phenotype of *nmr-2(ok3324)* animals, implicating a neuronal site of action (Figure 5A). Then we constructed a neuronal connectome of NMR-2-expressing neurons (Figure 5B) to uncover the specific neuron(s) through which it mediates the pathogen-induced immune response. Furthermore, to identify the specific neuron or neurons via which NMR-2 suppresses immunity, we studied animals with ablated individual NMR-2-expressing neurons: AVD, AVA, AVE, AVL, AVG, PVC, RIM, and RMD. Ablation of the AVD interneuron followed by pathogen exposure resulted in a survival phenotype similar to that of *nmr-2(ok3324)* animals, whereas ablation of other neurons, including AVA, AVE, AVL, AVG, PVC, RIM, or RMD, and exposure to pathogen produced survival comparable to wild-type animals (Figure 5C–D). Similar to *nmr-2(ok3324)*, AVD(-) animals showed no significant basal expression of immune genes but a significant upregulation of these genes upon pathogen exposure, as confirmed by qRT-PCR (Figure 5E–F). To determine whether *nmr-2* and AVD act in the same pathway, we generated an AVD(-);*nmr-2(ok3324)* animal and assessed their survival upon *S. aureus* exposure. These animals displayed a pathogen-resistant phenotype comparable to that of *nmr-2(ok3324)* animals (Figure 5G). To further determine whether *nmr-2* mediates pathogen resistance through the AVD neuron, we rescued AVD in *nmr-2(ok3324)* animals and exposed them to the pathogen. These animals exhibited a survival phenotype comparable to wild-type controls (Figure 5H), suggesting that NMR-2 regulates pathogen-induced immune response via the AVD interneuron.

**Figure 5.**
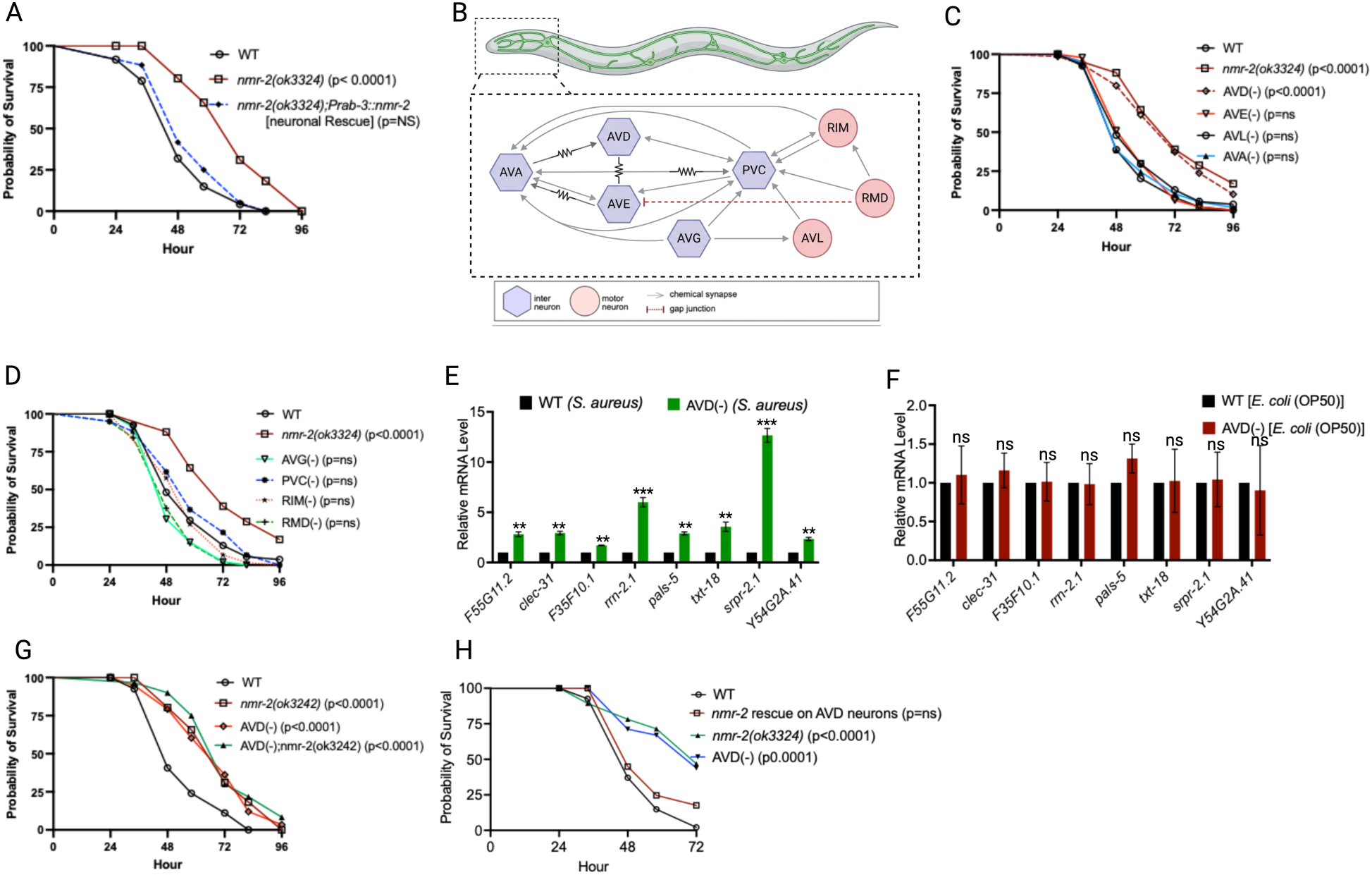
NMR-2 acts via the interneuron, AVD, to regulate pathogen-induced immunity A. WT, *nmr-2(ok3324), nmr-2(ok3324);Prab-3::nmr-2* (neuronal rescue) were exposed to *S. aureus* full lawn and scored for survival. B. Neuronal connectome of NMR-2-expressing neurons. C. WT, *nmr-2(ok3324),* AVD(-), AVE(-), AVL(-), and AVA(-) animals were exposed to *S. aureus* full lawn and scored for survival. D. WT, *nmr-2(ok3324),* AVG(-), PVC(-), RIM(-), and RMD(-) animals were exposed to *S. aureus* full lawn and scored for survival. E. qRT-PCR analysis of DAF-16 and PMK-1-dependent immune gene expression in WT and AVD(-) animals infected with *S. aureus*. Bars represent mean values, and error bars indicate the standard deviation (SD) from three independent experiments. Statistical significance is denoted as *p < 0.05, **p < 0.001, and ***p < 0.0001. F. qRT-PCR analysis of DAF-16 and PMK-1-dependent immune gene expression in WT and AVD(-) animals grown on *E. coli* (OP50). Bars represent mean values, and error bars indicate the standard deviation (SD) from three independent experiments. Statistical significance is denoted as *p < 0.05, **p < 0.001, and ***p < 0.0001. G. WT, *nmr-2(ok3324),* AVD(-), and AVD(-);*nmr-2(ok3324)* animals were exposed to *S. aureus* full lawn and scored for survival. H. WT, *nmr-2(ok3324),* AVD(-), AVD(-);*nmr-2(ok3324), nmr-2* rescue on AVD neurons animals were exposed to *S. aureus* full lawn and scored for survival.

### Sensory and Peripheral Neurons ASE, ASK, and AQR/PQR Signal Through AVD to Regulate Pathogen-Induced Immunity

Because NMR-2 regulates pathogen-induced immunity through the AVD interneuron (Figure 5), we examined AVD’s upstream connectivity. We built a neuronal connectome centered on the AVD interneuron, focusing on upstream sensory neurons known to regulate the immune response in *C. elegans* (Figure S5), and identified robust synaptic inputs from these sensory neurons: ASE, ASK, ASH, AQR, and PQR (Figure 6A). This connectivity reveals a defined sensory-to-interneuron circuit through which glutamatergic signaling may modulate systemic immune responses. Several sensory neurons that synapse onto AVD have previously been implicated in immune regulation in *C. elegans* ^7, 39, 40^. When animals lacking ASE, ASK, or AQR/PQR neurons were exposed to *S. aureus*, they exhibited pathogen resistance comparable to that of *nmr-2(ok3324)* mutants (Figure 6B–C). However, animals lacking ASH neurons also exhibited pathogen resistance that was less comparable to that of *nmr-2(ok3324)* animals (Figure 6D). In addition, ASE(–);*nmr-2(ok3324)*, ASK(–);*nmr-2(ok3324)*, and AQR/PQR(–);*nmr-2(ok3324)* animals displayed pathogen resistance comparable to *nmr-2(ok3324)* alone (Figure 6E–G). These results support a model in which individual or combined sensory neurons act upstream of NMR-2 and AVD within a neural circuit that regulates pathogen-induced immunity. Together, these findings demonstrate that NMDA receptor subunit NMR-2 functions through the AVD interneuron. This regulation likely involves communication with peripheral sensory neurons (ASE, ASK, and AQR/PQR) and modulates immune pathways dependent on DAF-16, PMK-1, and HLH-30 (Figure 7).

**Figure 6.**
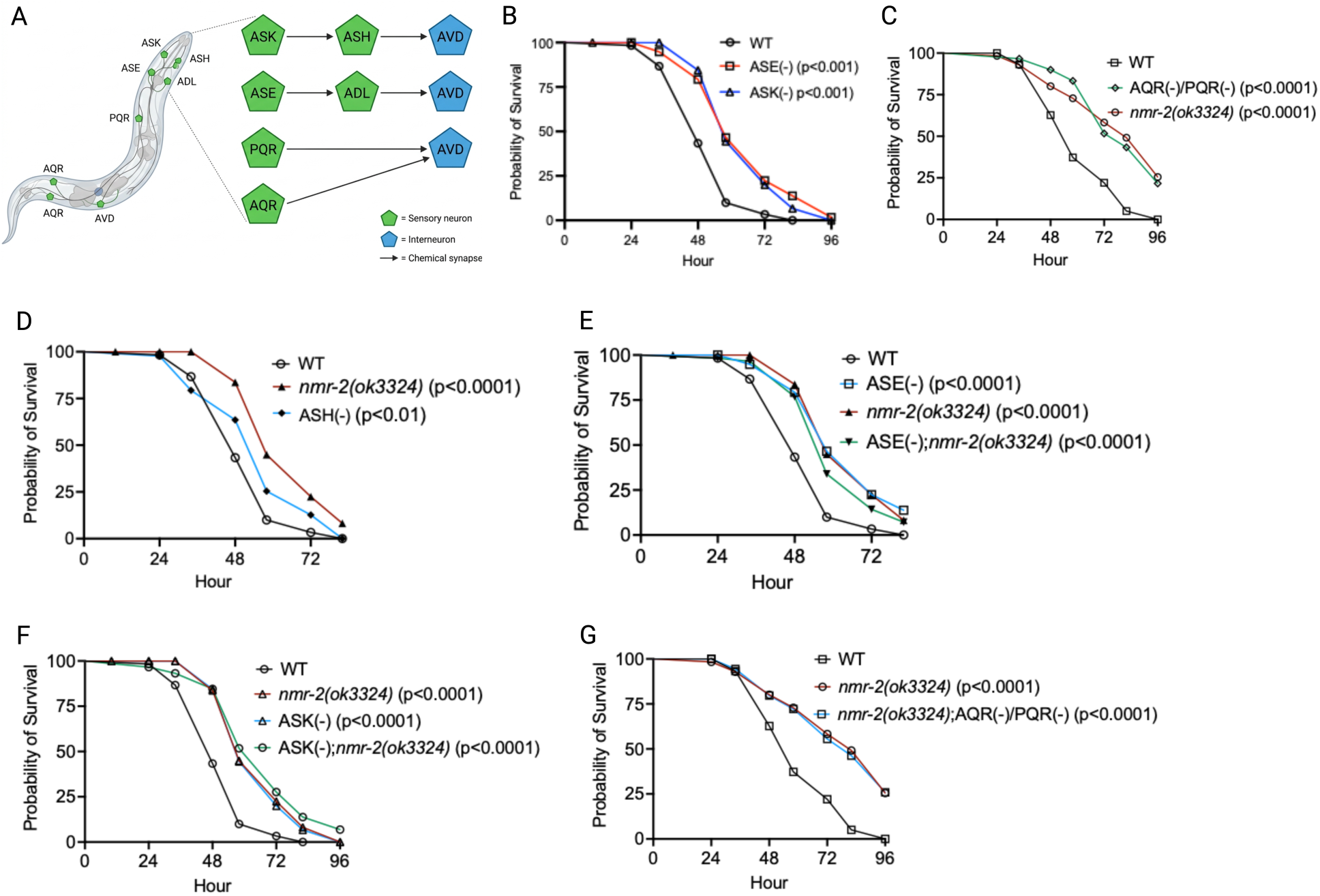
NMR-2 regulates pathogen-induced immunity through the AVD interneuron by signaling via sensory neurons ASE, ASK, and AQR/PQR. A. Sensory neuron connections to the NMR-2-expressing interneuron AVD. B. WT, *nmr-2(ok3324),* ASE(-), and ASK(-) animals were exposed to *S. aureus* full lawn and scored for survival. C. WT, *nmr-2(ok3324),* and AQR/PQR(-) animals were exposed to *S. aureus* full lawn and scored for survival. D. WT, *nmr-2(ok3324),* and ASH(-) animals were exposed to *S. aureus* full lawn and scored for survival. E. WT, *nmr-2(ok3324),* ASE(-), and ASE(-);*nmr-2(ok3324)* animals were exposed to *S. aureus* full lawn and scored for survival. F. WT, *nmr-2(ok3324),* ASK(-), and ASK(-);*nmr-2(ok3324)* animals were exposed to *S. aureus* full lawn and scored for survival. G. WT, *nmr-2(ok3324),* and AQR/PQR(-);*nmr-2(ok3324)* animals were exposed to *S. aureus* full lawn and scored for survival.

**Figure 7.**
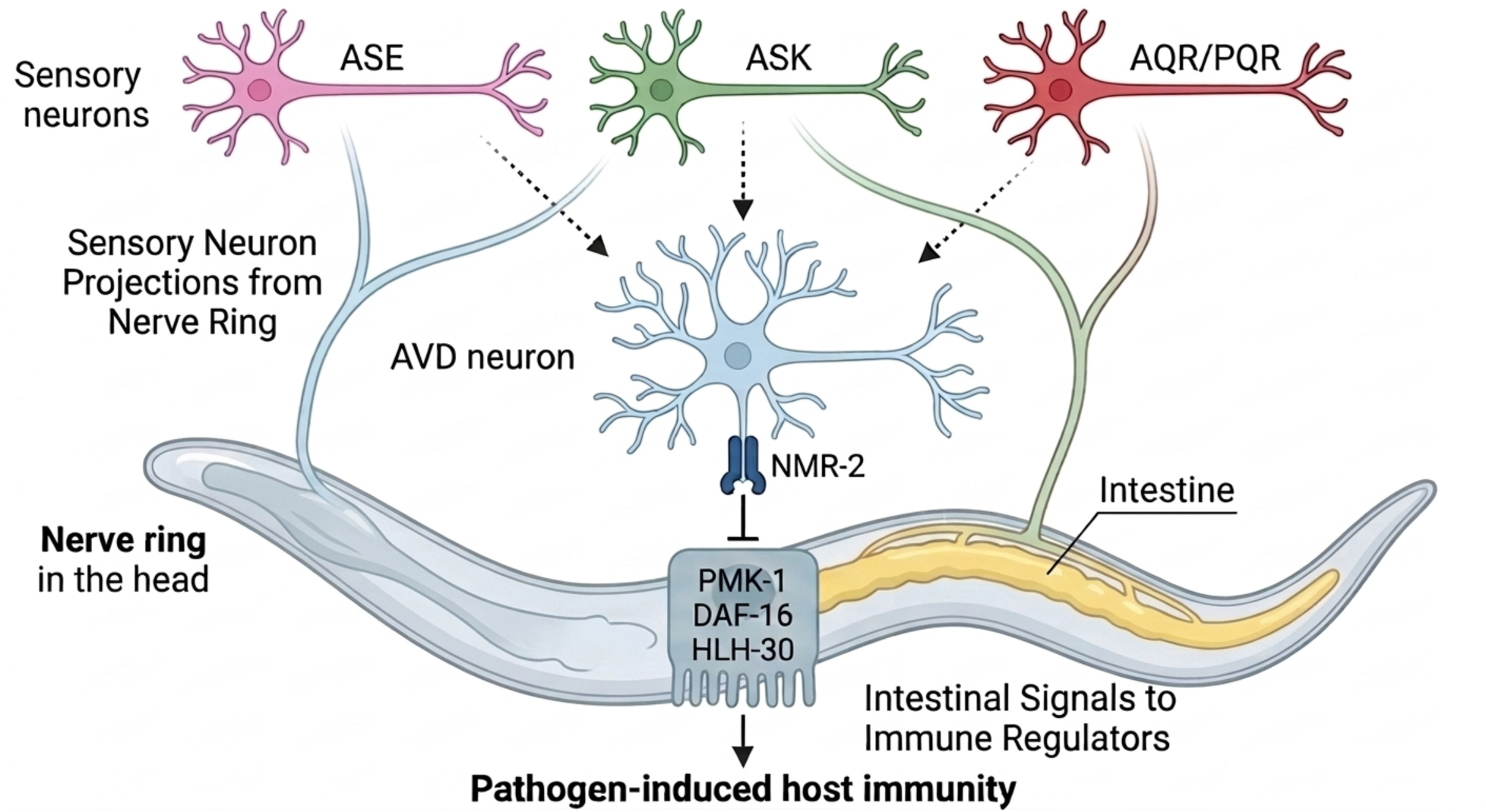
Neuronal NMR-2 regulates pathogen-induced immunity through the AVD interneuron via pathways dependent on PMK-1, DAF-16, and HLH-30. This immune function of NMR-2 in AVD may be mediated by input from upstream sensory neurons, including ASE, ASK, and AQR/PQR.

## Discussion

Glutamatergic signaling, a major neurotransmitter pathway, functions as the primary excitatory mechanism in the nervous system and is essential for learning, memory, and other cognitive functions across species, from humans to *C. elegans* ^17, 18, 22, 41, 42^. In *C. elegans*, this pathway is mediated in part by the NMDA receptor subunit NMR-2, which serves as a key component of glutamatergic transmission ^21^ and functions as a previously unrecognized regulator of pathogen-induced immunity, as demonstrated by the present work. NMR-2 modulates pathogen-induced immune responses through immune genes that are known to require the PMK-1/p38 MAPK, DAF-16/FOXO, and HLH-30/TFEB pathways. In *C. elegans*, NMDA receptors are heteromeric complexes composed of the essential subunit NMR-1, the homolog of mammalian GRIN1, and the modulatory subunit NMR-2, homologous to GRIN2 ^22^. Consistent with NMDA receptor architecture, NMR-1 is associated with co-agonist (glycine) binding, whereas NMR-2 contributes to glutamate binding ^17, 22^. This subunit organization may contribute to the differential phenotypes observed between *nmr-1* and *nmr-2* mutants, as loss of NMR-2 could selectively impair glutamatergic signaling without fully disrupting receptor function. In higher organisms, NMDA receptor signaling has been reported to exert both stimulatory and suppressive effects on immune responses in a context-dependent manner. In some settings, NMDA receptor activity promotes inflammatory signaling and cytokine production, whereas in others it contributes to immune suppression or maintenance of immune homeostasis, underscoring a conserved and bidirectional role in immune regulation ^43^.

In the current work, the pathogen-induced immune response is mediated through a specific interneuron, the AVD, and the upstream sensory neurons ASE, ASK, and AQR/PQR. Enhanced resistance in *nmr-2* mutants occurs without changes in bacterial colonization or avoidance behavior, indicating a direct effect on immune activation. The dissociation between survival and pathogen load is consistent with a tolerance-based mechanism in which survival is enhanced without a reduction in pathogen burden. Such tolerance mechanisms enable the host to withstand the detrimental effects of infection without necessarily reducing pathogen burden ^44, 45^. Consistent with this, previous studies in *C. elegans* have shown that increased survival can arise from enhanced tolerance rather than increased resistance^44, 45^.

The present work not only builds on a substantial body of literature demonstrating that the nervous system centrally orchestrates immune function in *C. elegans* ^7, 8, 10, 38, 46^ but also demonstrates a novel mechanism that controls inducible defense. Previous studies have shown that other neurotransmitters, such as serotonergic, dopaminergic, octopaminergic, and GABAergic, modulate innate immunity through specific neuronal circuits ^8, 9, 10, 14, 15^. For example, ASH octopaminergic neurons suppress immunity via GPCR signaling ^8, 14, 15^, dopaminergic neurons modulate defense through regulation of immune gene expression ^9^. Also, GABAergic function acts as an immunosuppressor via ASG neurons ^10^ , and serotonergic neurons act through MOD-1 to control resistance ^13^. Additionally, insulin-like peptides secreted by sensory neurons modulate DAF-16/FOXO activity and influence systemic immune readiness in *C. elegans* ^47^. Besides the fact that the role of glutamatergic signaling in host defense remains largely unexplored, none of the aforementioned neurotransmitter signaling pathways are known to regulate immunity specifically in response to pathogens rather than basal immunity. Consistent with the genetic analyses, pharmacological inhibition using dextromethorphan produced a similar survival phenotype in wild-type animals. However, because DXM is not fully selective for NMDA receptors, these findings should be considered supportive rather than definitive evidence for NMR-2-dependent glutamatergic signaling.

Here, we show that NMR-2-dependent glutamate signaling primarily modulates pathogen-induced immune responses without broadly altering basal immune gene expression, likely serving to minimize the energetic cost and collateral damage associated with chronic immune stimulation ^48, 49, 50^. Upon infection, functional loss of the inhibitory circuit uncovered in this study triggers a rapid and robust immune response, supporting a model in which NMR-2 limits pathogen-induced responses without evidence for constitutive basal immune suppression. This suggests that NMR-2 acts as a neural immune checkpoint, creating a tonic barrier that must be bypassed during infection to activate immune gene expression and ensure organismal protection, as observed in *C. elegans* ^15, 51^ and higher host organisms ^52, 53^. Such regulation may reflect a mechanism that balances pathogen-induced immune activation with the need to avoid inappropriate or excessive immune responses. Our data further demonstrated that the regulation of pathogen-induced immunity by NMR-2 acts via the AVD interneuron, which is traditionally known for its role in locomotor control and sensorimotor integration ^54, 55^. Although the precise mechanisms linking AVD neuronal activity to downstream immune pathways remain unclear, glutamatergic signaling may regulate neuroendocrine outputs that converge on immune regulators such as PMK-1, DAF-16, and HLH-30. Similar neuron-to-intestine signaling mechanisms have been described in *C. elegans*, where neuronal circuits coordinate systemic immune responses. This highlights the multifunctionality of interneurons in *C. elegans*, reinforcing a broader theme that individual neurons can regulate both behavior and physiology, including immunity ^7, 12, 14, 15^.

In vertebrates, glutamate receptors are expressed on various immune cells and are known to regulate key processes such as cytokine production, cell survival, inflammation, and innate immune responses ^56, 57, 58, 59, 60^. In addition to its role in immune cells, glutamatergic signaling has been implicated in coordinating systemic inflammatory and neuroimmune responses across different pathological conditions. Dysregulated glutamatergic signaling has been linked to sepsis, neuroinflammation, and autoimmune diseases, underscoring a potentially conserved role for glutamate in modulating systemic immune activation ^61, 62, 63, 64^. Our findings provide a framework for understanding neuroimmune mechanisms and pave the way for future studies on how glutamate interacts with other neuromodulators and environmental cues in regulating immunity across metazoans.

## Materials and Methods

### Bacterial strains

The bacterial strains used in this study are *Escherichia coli* OP50, *E. coli* HT115(DE3) ^65^, *P. aeruginosa* PA14, *P. aeruginosa* PA14-GFP ^66, 67^, and *Staphylococcus aureus* strain NCTCB325 ^68^. Gram-negative bacteria were grown in Luria-Bertani (LB) broth. *Staphylococcus aureus* strain NCTCB325 was grown in Tryptic Soy Agar prepared with nalidixic acid. All bacteria were grown at 37°C.

### *C. elegans* Strains and Growth Conditions

Hermaphrodite *C. elegans* (var. Bristol) Wild Type (WT) was used as a control unless otherwise indicated. *C. elegans* strains VC2623 *nmr-2(ok3324),* VM487 *nmr-1(ak4),* KP4 *glr-1 (n2461),* RB1808 *glr-2 (ok2342),* JIN1375 *hlh-30(tm1978),* CF1038 *daf-16(mu86)*, KU25 *pmk-1(km25),* GR2245 *skn-1(mg570)*, MGH171 *alxIs9* [*vha-6p::sid-1::SL2::GFP*], TU3401 *uIs69* [*pCFJ90 (myo-2p::mCherry) + unc-119p::sid-1*], JN1713 *peIs1713 [sra-6p::mCasp-1 + unc-122p::*mCherry (ASH neurons ablated strain), PR672 *che-1(p672)* (ASE neurons ablated strain), PS6025 *qrIs2 [sra-9::*mCasp1] (ASE neurons ablated strain) and AY101 acIs101 [F35E12.5p::GFP + rol-6(su1006)] were obtained from the Caenorhabditis Genetics Center (University of Minnesota, Minneapolis, MN) (https://cgc.umn.edu/). We also obtained FX06403 *glr-3(tm6403), glr-3(tm13827),* FX17095 *glr-4(tm31727),* FX03239 *glr-4(tm3239),* FX03506 *glr-5(tm3506),* FX19528 *mgl-1(tm1811),* FX31000 *mgl-2(tm8806),* FX00355 *mgl-2(tm355),* FX01766 *mgl-3(tm1766),* FX07159 *glr-8(tm7159),* FX07137 *glr-8(tm7137),* FX31703 *glr-7(tm13827)*, FX02877 *glr-7(tm2877),* FX02729 *glr-6(tm2729)* from the National Bioresource Project (NBRP), Japan (https://shigen.nig.ac.jp/c.elegans/mutants/index.xhtml). ASE(-);*nmr-2(ok3324)*, ASK(-); *nmr-2(ok3324)*, AQR/PQR;*nmr-2(ok3324)*, *daf-16(mu86); nmr-2(ok3324)*, *pmk-1(km25); nmr-2(ok3324), hlh-30(tm1978); nmr-2(ok3324),* were obtained by standard genetic crosses. Rescued strain *nmr-2(ok3324);Pnmr-2::nmr-2*, neuronal rescued strain *nmr-2(ok3324);Prab-3::nmr-2* were generated as described below. The neuron-ablated strains (carrying extrachromosomal arrays) targeting AVA, AVE, RIM, RMD, AVG, PVC, and AVD neurons were generated through services provided by SunyBiotech (https://www.sunybiotech.com/). The strains were crossed with the wild-type laboratory N2. All strains were grown at 20°C on nematode growth medium (NGM) plates seeded with *E. coli* OP50 as the food source ^65^ unless otherwise indicated. The recipe for the control NGM plates is: 3 g/l NaCl, 3 g/l peptone, 20 g/l agar, 5 μg/ml cholesterol, 1 mM MgSO4, 1 mM CaCl2, and 25 mM potassium phosphate buffer (pH 6.0). The NGM plates were without antibiotics except as indicated.

### RNA Interference (RNAi)

Gene knockdown was achieved by feeding animals with *E. coli* strain HT115(DE3) engineered to express double-stranded RNA (dsRNA) corresponding to the target gene ^69, 70^. RNAi was performed following previously described protocols ^8, 10, 37, 38^. Briefly, *E. coli* strains carrying the appropriate vectors were cultured overnight at 37°C in LB broth supplemented with ampicillin (100 μg/mL) and tetracycline (12.5 μg/mL). The cultures were then plated onto NGM plates containing ampicillin (100 μg/mL) and 3 mM isopropyl β-D-thiogalactoside (IPTG) (referred to as RNAi plates) and incubated at 37°C for 12–14 hours. Gravid adult worms were transferred onto the RNAi bacterial lawns and allowed to lay eggs for 2–3 hours before being removed. The resulting eggs were incubated at 20°C until they reached the young adult stage. This procedure was repeated for an additional generation, except for experiments involving ELT-2 RNAi, before the animals were used for assays. RNAi clones were obtained from the Ahringer RNAi library.

### *C. elegans* Survival Assay on Bacterial Pathogens

*Pseudomonas aeruginosa* was cultured in LB medium, while *Staphylococcus aureus* was grown in TSA medium supplemented with nalidixic acid (20 μg/mL). All bacterial cultures were incubated at 37°C with gentle shaking for 12 hours. For plate preparation, *P. aeruginosa* and *S. enterica* were seeded onto a modified NGM agar medium containing 0.35% peptone, and TSA was used for *S. aureus*. For partial lawn assays, 20 μL of the overnight bacterial culture was placed at the center of the agar plate without spreading. For full lawn assays, 20 μL of culture was evenly spread across the plate surface. No antibiotics were used for *P. aeruginosa* and *S. enterica* plates, while TSA plates for *S. aureus* contained nalidixic acid (20 μg/mL). Seeded plates were incubated at 37°C for 12 hours and then rested at room temperature for at least 1 hour before initiating infections. Twenty synchronized young adult *C. elegans* were transferred onto each infection plate. Three technical replicates (n = 60 animals) were set up per condition, and experiments were independently performed three times. Plates were incubated at 25°C, and survival was monitored every 12 hours for *P. aeruginosa* and *S. aureus*, and every 24 hours for *S. enterica*. Animals were considered dead if they failed to respond to a gentle touch with a worm pick or lacked pharyngeal pumping. Live animals were transferred daily to fresh pathogen lawns. All survival assays were repeated in three independent biological replicates.

### Bacterial lawn avoidance assay

Bacterial lawn avoidance assays were performed with 20 mL of *P. aeruginosa* PA14 and *S. aureus* NCTCB325 on 3.5-cm modified NGM agar plates (0.35% peptone) and (0.35 % TSA), respectively, which were cultured at 37°C overnight to have a partial lawn. The modified NGM plates were left to cool to room temperature for about 1 hour, and twenty young adult animals grown on *E. coli* OP50 were transferred to the center of each bacterial lawn after it. The number of animals on the bacterial lawns was counted at 12 and 24 hours after exposure.

### Pharyngeal pumping rate assay

Wild-type and *nmr-2(ok3324)* animals were synchronized by placing 20 gravid adult worms on NGM plates seeded with *E. coli* OP50 and allowing them to lay eggs for 60 min at 20°C. The gravid adult worms were then removed, and the eggs were allowed to hatch and grow at 20°C until they reached the young adult stage. The synchronized worms were transferred to NGM plates fully seeded with *P. aeruginosa* for 24 hours at 25°C. Worms were observed under the microscope with a focus on the pharynx. The number of contractions of the pharyngeal bulb was counted over 60 s. Counting was conducted in triplicate and averaged to obtain pumping rates.

### Defecation rate assay

Wild-type and *nmr-2(ok3324)* animals were synchronized by placing 20 gravid adult worms on NGM plates seeded with *E. coli* OP50 and allowing them to lay eggs for 60 min at 20°C. The gravid adult worms were then removed, and the eggs were allowed to hatch and grow at 20°C until they reached the young adult stage. The synchronized worms were transferred to NGM plates fully seeded with *P. aeruginosa* for 24 hours at 25°C. Worms were observed under a microscope at room temperature. For each worm, an average of 10 intervals between two defecation cycles was measured. The defecation cycle was identified as a peristaltic contraction beginning at the posterior body of the animal and propagating to the anterior part of the animal followed by feces expulsion.

### Brood Size Assay

The brood size assay was done following the earlier described methods ^71, 72^. Ten L4 animals from egg-synchronized populations were transferred to individual NGM plates (seeded with *E. coli* OP50) (described above) and incubated at 20°C. The animals were transferred to fresh plates every 24 hours. The progenies were counted and removed every day.

### *C. elegans* Longevity Assays

Longevity assays were performed on NGM plates containing live, UV-killed *E. coli* strains HT115 or OP50 as described earlier ^8, 73, 74, 75, 76^. Animals were scored as alive, dead, or gone each day. Animals that failed to display touch-provoked or pharyngeal movement were scored as dead. Experimental groups contained 60 to 100 animals, and the experiments were performed in triplicate. The assays were performed at 20°C.

### Intestinal Bacterial Loads Visualization and Quantification

Intestinal bacterial loads were visualized and quantified as described earlier ^8, 73^. Briefly, *P. aeruginosa*-GFP lawns were prepared as described above. The plates were cooled to ambient temperature for at least an hour before seeding with young gravid adult hermaphroditic animals, and the setup was placed at 25^°^C for 24 hours. The animals were transferred from *P. aeruginosa*-GFP plates to the center of fresh *E. coli* plates for 10 min to eliminate *P. aeruginosa*-GFP from their body. The step was repeated two times more to further eliminate external *P. aeruginosa*-GFP left from earlier steps. Subsequently, ten animals were collected and used for fluorescence imaging to visualize the bacterial load, while another ten were transferred into 100 µL of PBS plus 0.01% Triton X-100 and ground. Serial dilutions of the lysates (10^1^-10^10^) were seeded onto LB plates containing 50 µg/mL of kanamycin to select for *P. aeruginosa*-GFP cells and grown overnight at 37 °C. Single colonies were counted the next day and represented as the number of bacterial cells or CFU per animal.

### Glutamate and Dextromethorphan Hydrobromide Treatment

Dextromethorphan hydrobromide (DXM) stock solution (40 mg/mL) was prepared by dissolving 400 mg of the compound in 10 mL of 100% ethanol. The stock solution was added to SB medium to achieve a final concentration of 40 µg/mL. Control plates were prepared by adding an equivalent volume of 100% ethanol to the SB medium. WT and *nmr-2(ok3324)* animals were maintained on the respective SB plates from egg stage to young adulthood, after which they were transferred to *S. aureus* lawns for survival assays as described above.

### Fluorescence Imaging

Fluorescence imaging was carried out as described previously ^72^. Briefly, animals were anesthetized using an M9 salt solution containing 50 mM sodium azide and mounted onto 2% agar pads. The animals were then visualized for bacterial load using a Leica M165 FC fluorescence stereomicroscope. The diameter of the intestinal lumen was measured using Fiji-ImageJ software. At least 10 animals were used for each condition.

### Fluorescent Reporter Assays (AY101 Strain)

To evaluate basal and pathogen-induced immune gene expression at single-animal resolution, we employed the transgenic reporter strain AY101 [*pirg-5p::GFP + rol-6(su1006)*], in which GFP is driven by the promoter of the immune effector gene *irg-5*. AY101 animals were maintained under standard conditions on *E. coli* OP50 lawns at 20 °C prior to experiments. For assessment of basal immune activity, individual day 1 adult animals were examined on E. coli plates using a fluorescence microscope. GFP expression was qualitatively classified as either “bright” (strong intestinal fluorescence) or “low/undetectable” (faint or absent fluorescence). For infection assays, synchronized day 1 adults were transferred to full lawns of Staphylococcus aureus strain NCTC8325 and monitored for survival as described above. Animals were stratified based on their basal GFP levels (bright vs. low/undetectable) before infection, and survival curves were generated for each group.

### RNA Sequencing and Bioinformatic Analyses

Approximately 40 gravid wild-type and *nmr-2(ok3324)* animals were placed on 10-cm NGM plates seeded with *E. coli* OP50 for 3 hours to obtain synchronized populations. The animals were then allowed to develop to the L4 larval stage at 20°C. A separate group of animals was infected with *S. aureus* for 4 hours as described above. Following treatment, animals were washed off the plates with M9 buffer, flash-frozen in QIAzol using an ethanol/dry ice bath, and stored at –80°C until RNA extraction. Total RNA was isolated using the RNeasy Plus Universal Kit (Qiagen, Netherlands). Residual genomic DNA was removed with the TURBO DNase kit (Life Technologies, Carlsbad, CA). For cDNA synthesis, 6 μg of total RNA was reverse-transcribed using random primers and the High-Capacity cDNA Reverse Transcription Kit (Applied Biosystems, Foster City, CA).

The library construction and RNA sequencing in Illumina NovaSeq 6000 platform were done following the method described by ^77^ and ^78^. Pair-end reads of 150 bp were obtained for subsequent data analysis. The RNA sequence data were analyzed using a workflow constructed for Galaxy (https://usegalaxy.org) as described ^79^ and were validated using Lasergene DNA Star software. The RNA reads were aligned to the *C. elegans* genome (WS271) using the aligner STAR. Counts were normalized for sequencing depth and RNA composition across all samples. Differential gene expression analysis was then performed on normalized samples. Genes exhibiting at least two-fold change were considered differentially expressed. The differentially expressed genes were subjected to SimpleMine tools from wormbase (https://www.wormbase.org/tools/mine/simplemine.cgi) to generate information such as wormBase ID and gene name, which are employed for further analyses. Gene ontology analysis was performed using the WormBase IDs in DAVID Bioinformatics Database (https://david.ncifcrf.gov) ^80^ and validated using a *C. elegans* data enrichment analysis tool (https://wormbase.org/tools/enrichment/tea/tea.cgi). Several genes in our dataset were listed from the WormBase enrichment analysis tool as valid but unannotated. As subsequent *C. elegans* genome releases may update gene annotations and Gene Ontology (GO) terms, minor differences from the current analysis are expected. To gain further insight into immune regulation, we compared the upregulated genes to known targets of key immune pathways, including PMK-1, DAF-16, ELT-2, SKN-1, and HLH-30 obtained from Worm Exp version 1 (http://wormexp.zoologie.uni-kiel.de/wormexp/) ^81^ using the transcription factor target category. Enrichment factor and corresponding P value were calculated using the Overlap Statistics Calculator tool http://nemates.org/MA/progs/overlap_stats.html. The Venn diagrams were obtained using the web tool InteractiVenn (http://www.interactivenn.net) ^82^ and a bioinformatics and evolutionary genomics tool (http://bioinformatics.psb.ugent.be/webtools/Venn/). Neuron wiring was done using the database of synaptic connectivity of *C. elegans* for computation ^83^ http://ims.dse.ibaraki.ac.jp/ccep-tool/. RNA-seq datasets for *octr-1(ok371)* worms were downloaded from the NCBI GEO database (GSE252054) ^16^ in FASTQ format and subsequently reanalyzed as described above for the current work. The heat map was generated from selected genes using Morpheus, an online visualization tool developed by the Broad Institute https://software.broadinstitute.org/morpheus.

### RNA Isolation and Quantitative Reverse Transcription-PCR (qRT-PCR)

Approximately 40 gravid WT, *nmr-2(ok3324),* AVD(-) animals were placed on 10-cm NGM plates seeded with *E. coli* OP50 for 3 hours to obtain synchronized populations. The animals were then allowed to develop to the L4 larval stage at 20°C. A separate group of animals was infected with *S. aureus* for 4 hours as described above. Following standard protocol, total RNA extraction was done following the protocol as described above. qRT-PCR was conducted using the Applied Biosystems One-Step Real-time PCR protocol using SYBR Green fluorescence (Applied Biosystems) on an Applied Biosystems 7900HT real-time PCR machine in 96-well-plate format. Twenty-five microliter reactions were analyzed as outlined by the manufacturer (Applied Biosystems). The relative fold changes of the transcripts were calculated using the comparative *CT*(2^−ΔΔ*CT*^) method and normalized to pan-actin (*act-1, -3, -4*). The cycle thresholds of the amplification were determined using StepOnePlus™ Real-Time PCR System Software v2.3 (Applied Biosystems). All samples were run in triplicate. The primer sequences were available upon request and presented in Supplementary Table 5.

### Generation of transgenic *C. elegans*

To generate *nmr-2* rescue animals, the DNA *nmr-2* alongside its promoter was amplified from the genomic DNA of Bristol N2 *C. elegans* adult worms. Plasmid pPD95_77_*Pnmr-2-_nmr-2* was constructed by linearization of plasmid pPD95_77 using SacI and EagI restriction enzymes. The amplified *Pnmr-2::nmr-2* DNA was cloned behind its native promoter in the plasmid pPD95_77, between SacI and EagI sites. For the neuronal rescue strain, the plasmid pPD95_77_*Prab-3_nmr-2* was constructed by cloning amplified *npr-15* DNA into SphI and ApaI digested pPD95_77_*Prab-3* under the promoter of *rab-3*. The constructs were purified and sequenced. Young adult hermaphrodite *nmr-2(ok3324)* and WT *C. elegans* were transformed by microinjection of plasmids into the gonad as described ^84, 85^. Briefly, a mixture containing pPD95_77_*Pnmr-2_nmr-2* (40 ng/μl) and *Pmyo-3*::mCherry (5 ng/μl) that drives the expression of mCherry to the muscle as a transformation marker was injected into the animals. For the neuronal rescue, a mixture containing pPD95_77_*Prab-3_nmr-2* plasmids (40 ng/μl) and *Pmyo-3*::mCherry (5 ng/μl) that drives the expression of mCherry as a transformation marker was injected into the animals.

### Neuronal Ablation Constructs and Plasmid Design

Neuronal ablation was achieved using both single-caspase and intersectional split-caspase strategies. For split-caspase systems, complementary fragments of CED-3 (p17 and p15) were expressed under distinct promoters to induce apoptosis selectively in overlapping neuronal populations. Constructs were generated by replacing promoter sequences in standard vectors pPD95.77 using SphI and NheI restriction sites, with all plasmids containing the unc-54 3′UTR for transcript stability.

The promoter-caspase combinations used were as follows: AVA, *Pflp-18*::CZ*-ced*-3(p17) + *Pgpa-14::ced-3*(p15)-NZ (split-caspase); AVE, *Popt-3::ced-3* (single-caspase);RIM, *Pgcy-13*::ICE (human caspase-1);RMD, PC42D4.1::ICE (human caspase-1);AVG, *Pnmr-1*::CZ-*ced-3(p17)* + *Pflp-7::ced-3*(p15)-NZ (split-caspase);PVC, *Pnmr-1::*CZ*-ced-3*(p17) + *Pflp-20::ced-3*(p15)-NZ (split-caspase);AVD, *Pcfi-1::*CZ*-ced-3*(p17) + *Pprx-3a::ced-3*(p15)-NZ (split-caspase). All constructs were verified by Sanger sequencing prior to transgenesis. Generation of Transgenic Lines

Transgenic *C. elegans* were generated by microinjection of plasmid DNA into the gonads of young adult N2 hermaphrodites. Injection mixtures contained 25 ng/µL of each construct and 50 ng/µL pRF4 [*rol-6(su1006)*] as a co-injection marker. For the RMD ablation construct, 5 ng/µL plasmid DNA was used with 45 ng/µL pSL1190 carrier DNA to minimize toxicity. F1 Roller progeny were selected to establish stable extrachromosomal array lines. Multiple independent lines were generated and maintained under standard conditions to account for variability and mosaicism.

### AVD-Specific Rescue of *nmr-2*

Cell-specific rescue of *nmr-2* in AVD neurons was achieved using an intersectional split-intein/GAL4-UAS system. Three plasmids were co-injected into the *nmr-2(ok3324)* background: (i) *Pcfi-1*::NLS–*gp41* C-intein–VP64, which drives expression in AVD interneurons, (ii) Pprx-3::NLS–GAL4–*gp41* N-intein, providing complementary intein fragments, and (iii) 14xUAS::nmr-2::let-858 3′UTR, enabling reconstitution and expression of *nmr-2* specifically in overlapping cells. Plasmids were injected at ∼10 ng/µL each along with a co-injection marker (*Pmyo-3::*mCherry, 5 ng/µL), and stable extrachromosomal arrays were generated following standard microinjection procedures. Transgenic lines were identified based on marker expression and maintained for subsequent phenotypic and survival analyses.

### Quantification and Statistical Analysis

Statistical analysis was performed using Prism 8 (version 8.1.2; GraphPad). For non-survival assays, error bars represent the standard deviation (SD) from at least three independent experiments. Statistical significance was assessed using two-sample t-tests where appropriate, with significance defined as p < 0.05. In figures, asterisks denote significance as follows: ns, not significant; *, p < 0.05; **, p < 0.001; ***, p < 0.0001, compared with the appropriate controls. Survival data were analyzed using the Kaplan–Meier method, and statistical differences between curves were evaluated using the log-rank test. Consistent with standard practice in C. elegans survival assays, survival curves are presented without error bars, as variability across replicate experiments is incorporated into the statistical analysis rather than displayed graphically. All experiments were performed at least three times.

## ACKNOWLEDGMENTS

This work was fully supported by NIH grants GM0709077 and AI117911 (to A.A.). Most strains used in this study were obtained from the Caenorhabditis Genetics Center (CGC), which is funded by the NIH Office of Research Infrastructure Programs (P40 OD010440) and the National BioResource Project (NBRP) of Japan.

## AUTHOR CONTRIBUTIONS

B.O. and A.A. conceived and designed the experiments. B.O. and JL performed the experiments. B.O. and A.A. analyzed the data and wrote the paper.

## DECLARATION OF INTERESTS

The authors declare no competing interests.

## DATA AVAILABILITY

The RNA-seq datasets generated in this study have been deposited in the NCBI Gene Expression Omnibus (GEO) under accession number GSE261802.

---

**Figure S1.**
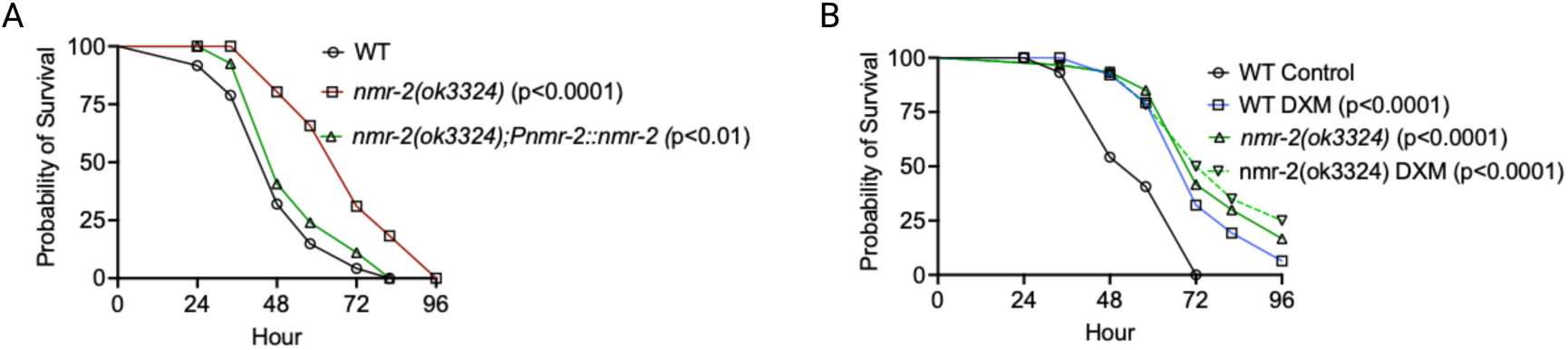
Role of nmr-2 in survival during S. aureus infection and response to dextromethorphan (DXM) A. WT, *nmr-2(ok3324),* and *nmr-2(ok3324);Pnmr-2::nmr-2* were exposed to *S. aureus* full lawn and scored for survival. B. WT and *nmr-2(ok3324)* treated with dextromethorphan compound were exposed to *S. aureus* full lawn and scored for survival.

**Figure S2.**
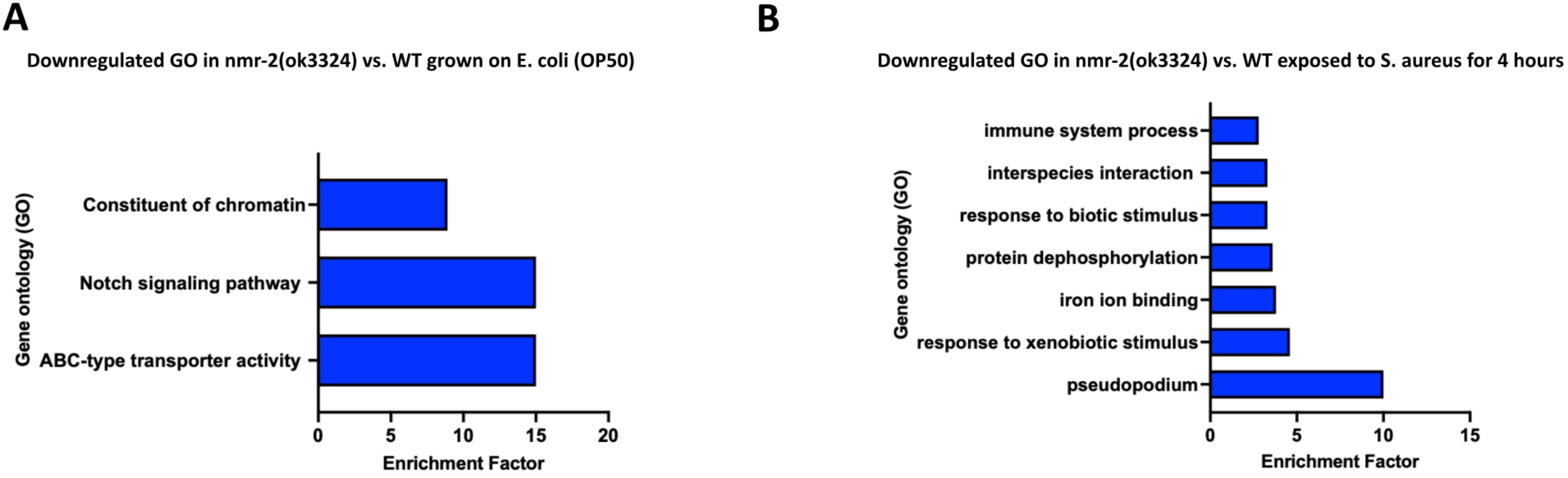
Gene ontology analysis of downregulated genes in *nmr-2(ok3324)* compared to wild-type animals grown on *E. coli* (OP50) and exposed to *S. aureus* for four hours. A. Gene ontology analysis of downregulated genes in *nmr-2(ok3324)* versus WT animals grown on *E. coli* (OP50). The results were filtered to include only significantly enriched terms with a q-value < 0.1, with minor modifications made by merging similar Gene Ontology terms where appropriate. B. Gene ontology analysis of downregulated genes in *nmr-2(ok3324)* versus WT animals exposed to *S. aureus* for four hours. The results were filtered to include only significantly enriched terms with a q-value < 0.1, with minor modifications made by merging similar Gene Ontology terms where appropriate.

**Figure S3.**
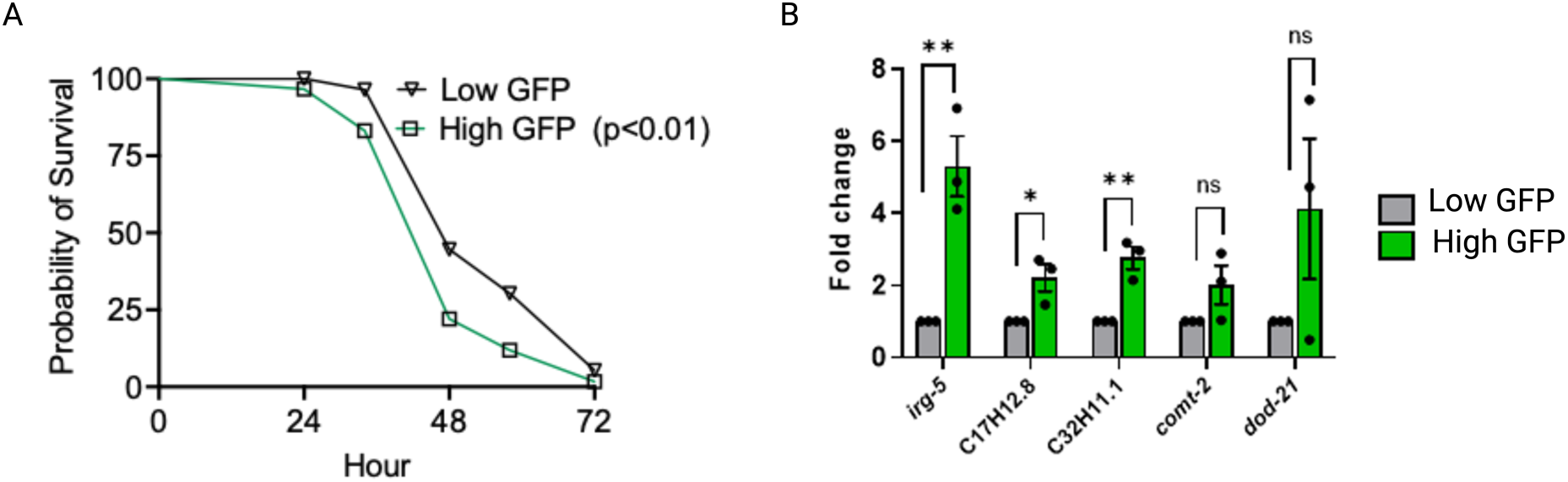
Differential survival and immune responses of low and high GFP AY101 animals A. Low and high Green Fluorescent Protein (GFP) AY101 animals were exposed to *S. aureus* full lawn and scored for survival. B. Quantitative reverse transcription-PCR (qRT-PCR) analysis was performed to examine Low and high Green Fluorescent Protein (GFP) AY101 animals’ immune gene expression. Bars represent mean values, and error bars indicate the standard deviation (SD) from three independent experiments. Statistical significance is denoted as *p < 0.05, **p < 0.001, and ***p < 0.0001.

**Figure S4.**
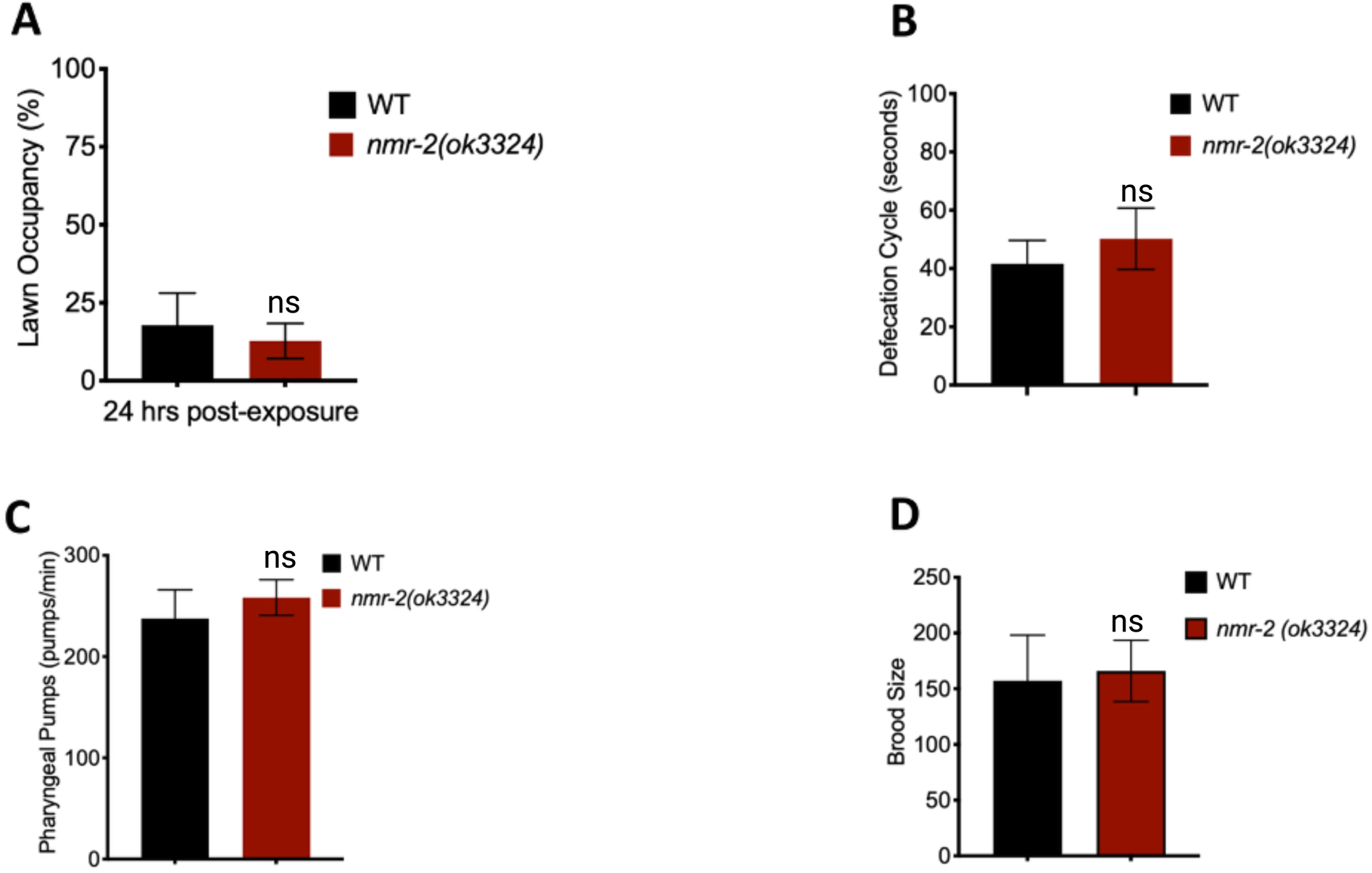
The enhanced pathogen resistance phenotype observed in *nmr-2* mutants is independent of pathogen avoidance, defecation rate, pharyngeal pumping, and brood size. A. Lawn Occupancy of WT and *nmr-2(ok3324)*. WT vs *nmr-2(ok3324)*, P = NS. B. Defecation cycle of WT and *nmr-2(ok3324)*. WT vs *nmr-2(ok3324)*, P = NS. C. Pharyngeal pumping of WT and *nmr-2(ok3324)*. WT vs *nmr-2(ok3324)*, P = NS. D. Brood size of WT and *nmr-2(ok3324)*. WT vs *nmr-2(ok3324)*, P = NS.

**Figure S5.**
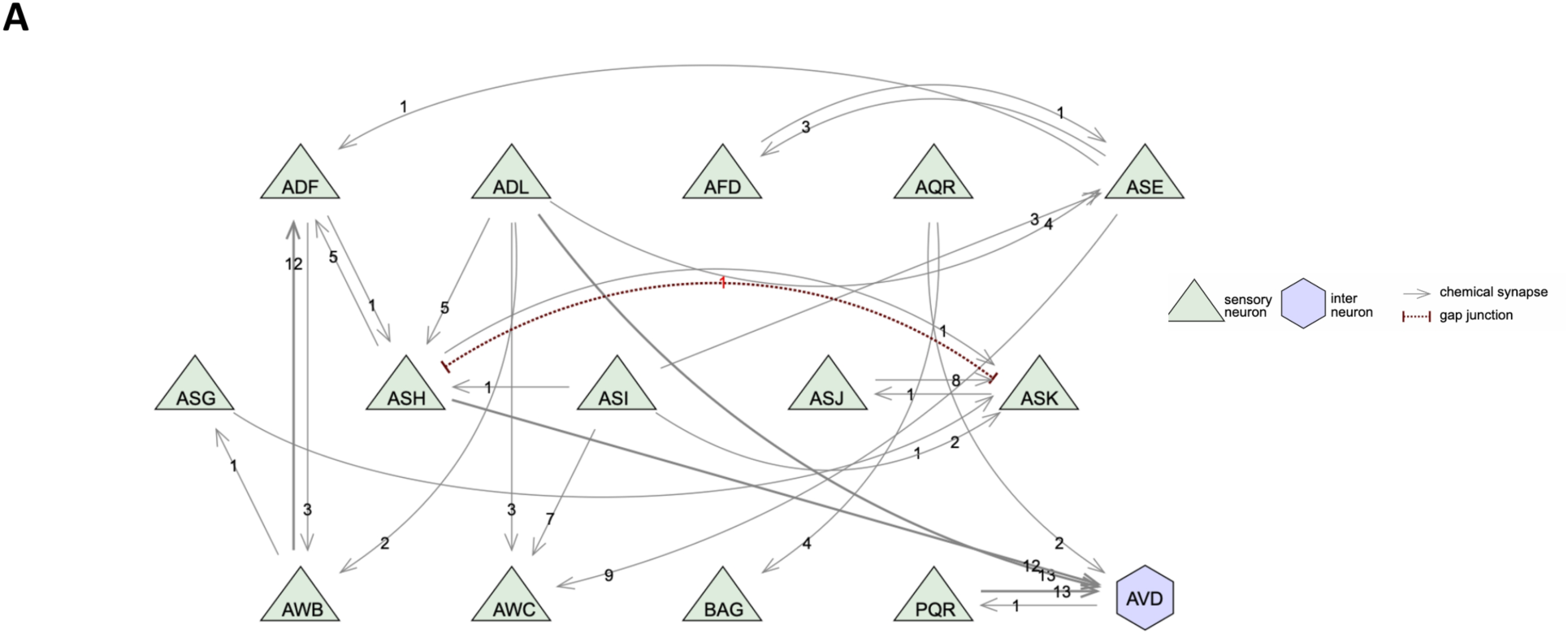
Connectome of immunologically relevant upstream neurons projecting to the AVD interneuron

## Notes

### Competing Interest Statement

The authors have declared no competing interest.

